# Unlocking the potential of historical abundance datasets to study biomass change in flying insects

**DOI:** 10.1101/695635

**Authors:** Rebecca S. Kinsella, Chris D. Thomas, Terry J. Crawford, Jane K. Hill, Peter J. Mayhew, Callum J. Macgregor

**Affiliations:** Department of Biology, University of York, Wentworth Way, York, YO10 5DD; Leverhulme Centre for Anthropocene Biodiversity, University of York, Wentworth Way, York, YO10 5DD; Energy and Environment Institute, University of Hull, Cottingham Road, Hull, HU6 7RX

**Keywords:** Biodiversity decline, body mass, forewing length, Lepidoptera, moths, predictive model

## Abstract

1. Trends in insect abundance are well-established in some datasets, but far less is known about how abundance measures translate into biomass trends. Moths (Lepidoptera) provide particularly good opportunities to study trends and drivers of biomass change at large spatial and temporal scales, given the existence of long-term abundance datasets. However, data on the body masses of moths are required for these analyses, but such data do not currently exist.
2. To address this data gap, we collected empirical data in 2018 on the forewing length and dry mass of field-sampled moths, and used these to train and test a statistical model that predicts the body mass of moth species from their forewing lengths (with refined parameters for Crambidae, Erebidae, Geometridae and Noctuidae).
3. Modelled biomass was positively correlated, with high explanatory power, with measured biomass of moth species (R^2^ = 0.886 ± 0.0006, across 10,000 bootstrapped replicates) and of mixed-species samples of moths (R^2^ = 0.873 ± 0.0003), showing that it is possible to predict biomass to an informative level of accuracy, and prediction error was smaller with larger sample sizes.
4. Our model allows biomass to be estimated for historical moth abundance datasets, and so our approach will create opportunities to investigate trends and drivers of insect biomass change over long timescales and broad geographic regions.

## Background

Multiple recent studies have reported that insect biomass, abundance and diversity are in decline over recent decades (Conrad *et al*., 2006; Hallmann *et al*., 2017, 2019; Lister & Garcia, 2018; Harris et al., 2019; Seibold *et al*., 2019; van Strien *et al*., 2019; Wepprich *et* al., 2019; Bell *et al*., 2020; Roth *et al*., 2020; Salcido *et al*., 2020), but with substantial spatial, temporal and taxonomic variation in the existence and strength of such declines (Shortall *et al*., 2009; Macgregor *et al*., 2019b; Outhwaite *et al*., 2020). The reasons for such variation are not yet known, and it is therefore possible that declines in insect biomass are not always symptomatic of equivalent declines in abundance, or *vice versa*. Biomass could remain stable even in the face of declining abundance if communities became increasingly comprised of larger-bodied species (Pöyry *et al*., 2017). Likewise, changes in community-level biomass could be attributable to changes in community composition, even in the absence of an overall abundance change. This might occur if insect communities became more biased towards larger-or smaller-bodied species, e.g. through size-bias in strength of selection for or against particular traits (Coulthard *et al*., 2019), such as faster or slower life-histories, degree of habitat specialization (Mattila *et al*., 2011; Davis *et al*., 2013), or strength of attraction to artificial light at night (van Langevelde *et al*., 2011). However, the dynamics of biomass and of biomass change, as well as the relationships between biomass, abundance, species richness, and community composition, remain poorly understood at large spatial and temporal scales because of a lack of suitable data on insect biomass (Macgregor *et al*., 2019b).

Opportunities to investigate changes over time and space in insect communities are provided by several large-scale, long-term abundance datasets for moths (Lepidoptera) in the UK, including the Rothamsted Insect Survey (RIS; Storkey *et al*., 2016), the National Moth Recording Scheme (NMRS; Fox *et al*., 2011), and the Garden Moth Scheme (GMS; Bates *et al*., 2014; Wilson *et al*., 2018), and elsewhere (e.g. Groenendijk & Ellis, 2011; Valtonen *et al*., 2014, 2017). Two-thirds of British species of macro-moths show negative abundance trends in the long-term (Conrad *et al*., 2006), with similar patterns observed elsewhere in Europe (e.g. Groenendijk & Ellis, 2011; Valtonen *et al*., 2017). The potential drivers of these declines are diverse (Fox, 2013), and likely to include habitat loss and fragmentation, agricultural intensification and associated agrochemical use, increased prevalence of artificial light at night and other factors associated with urbanisation, and climate change (Wickramasinghe *et al*., 2004; Morecroft *et al*., 2009; Fox, 2013; Fox *et al*., 2014; Gilburn *et al*., 2015; van Langevelde *et al*., 2018). Moths contribute to important ecosystem functions, including nocturnal pollination (Macgregor *et al*., 2015, 2019a) and energy transfer from producers to higher-level consumers (e.g. Franklin *et al*., 2003; Hooks *et al*., 2003; Singer *et al*., 2012). Thus, moths can be important to the conservation of their predators, such as bats (Vaughan, 1997; Threlfall *et al*., 2012) and some birds (Sierro *et al*., 2001; Denerley *et al*., 2018). In transferring energy, the quantity of vegetation consumed by caterpillars and the biomass of insects available to predators may be functionally important determinants of ecosystem processes (Brose *et al*., 2005). Similarly, the body size of individual species can play a substantial role in structuring networks of interspecific interactions (Woodward *et al*., 2005). All of these factors make moths a valuable taxon in which to study long-term biomass change at the community level, but biomass data are currently lacking for these analyses.

Existing long-term moth population and distribution datasets are potentially a very valuable resource for understanding biomass changes, but these datasets record abundance, not measurements of body mass or size, and in most cases do not retain specimens (preventing biomass information from being obtained retrospectively). To address questions of biomass change using these abundance datasets requires reliable body mass data for all species, but such empirical data are currently available for only a limited set of species (García-Barros, 2015). An alternative approach is to use empirical data from a subset of species to model the expected body mass of all species, using some other, more readily-available, trait. Such models have previously been formulated to predict the body mass of moths and other invertebrates from their body length (Sage, 1982; Sample *et al*., 1993; Sabo et al., 2002; Höfer & Ott, 2009) and variants thereof (García-Barros, 2015), chosen because it is easily measurable from museum specimens (García-Barros, 2015). However, for moths, body length data are not widely available and in any case may be influenced to a greater degree by contraction in dried specimens than other traits (García-Barros, 2015). The only morphological trait for which existing data on many species are readily available is forewing length: for example, an expected range of forewing lengths is included for all British species of macro-moths, and most British species of micro-moths, in standard field guides (Sterling & Parsons, 2012; Waring & Townsend, 2017), and it may therefore be possible to predict body mass based on forewing length (Miller, 1997). The existence of substantial interfamilial variation in body plan (e.g. between Saturniidae and Sphingidae; Janzen, 1984) may provide opportunities to use taxonomy to fine-tune models, but no previous model has included any refinement based on taxonomic relationships between moths.

In this study, we develop a statistical model to estimate the body mass of individual moths from their forewing length, and hence quantify the biomass of samples of moths for which species abundances only have been recorded. We have four aims: (i) collection of empirical data (during 2018 on the University of York campus, UK) to establish the relationship between forewing length and body mass in moths; (ii) construction of a predictive model for estimating body mass from species identity and associated forewing length, (iii) testing the accuracy of this model’s predictions and how accuracy changes with increasing moth abundance, and (iv) using existing data on forewing lengths to predict the body masses of all British macro-moths, thus providing a resource to users of moth population data and to comparative biologists.

## Materials and methods

### Field sampling, identification and measurement of moths

We sampled moths at three sites (Fig. S1) on the University of York campus, UK (53°56’41” N 1°2’2” W; vice-county 61, South-east Yorkshire) between 11th June and 20th July 2018 (Appendix S1.1). Moths were collected using Heath-style moth traps (Heath, 1965), each operating a 15 W actinic fluorescent tube and powered by a 12 V battery (Anglian Lepidopterist Supplies, Hindolveston, UK). Moths were euthanized and returned to the laboratory for identification and measurement. Moths were identified to species-level where possible using standard field guides (Sterling & Parsons, 2012; Waring & Townsend, 2017). Where species-level identification would have required dissection of the genitalia, identification was made to aggregate level (e.g. Common Rustic agg. *Mesapamea secalis/didyma*). After identification, moths were allowed to air-dry at room temperature for a minimum of one week, which was sufficient for the dry body mass of even the largest individuals to stabilize (Appendix S1.2, Fig. S2). After drying, we measured the forewing length and dry mass of each moth. Forewing length was measured from wing base to wing-tip, using calipers and a ruler, to the nearest 1 mm. Dry mass was measured using an A&D HR-202 balance (A&D Instruments Ltd., Abingdon, UK), to the nearest 0.01 mg. Measurements were precise to within ± 6 % of the true value (Appendix S1.2).

### Modelling forewing length – body mass relationship from empirical data

To investigate the relationship between forewing length (mm) and body mass (mg) in moths, we constructed generalized linear mixed-effects models (GLMMs) using our 2018 field data, with species as a random effect, and body mass explained by the interaction between forewing length and taxonomic family. We selected between three candidate model structures by comparison of Bayesian Information Criterion (BIC) scores: (i) linear predictor (i.e. ln(body mass) ∼ wing length × family); (ii) non-linear predictor (i.e. ln(body mass) ∼ ln(wing length) × family); and (iii) segmented predictor (as for model i, but permitting the slope of the model to change once as forewing length increases). Finally, we tested the significance of independent variables, including the interaction between wing length and family, using Likelihood Ratio Tests.

To reduce the risk of our predictive model overfitting for families represented by only a few species in our dataset (and therefore to allow accurate predictions of body mass to be made), we refitted this model with a simplified family variable, in which seven families represented by fewer than five species in our dataset were grouped together as ‘other’ (effectively reducing the family variable from 11 categories to 5). The four retained families (each with ≥ 5 species sampled) were Crambidae, Erebidae, Geometridae and Noctuidae, allowing the predictive model’s parameters to be refined for these families, whilst also making overall predictions for all other families. We fitted a GLMM to the dataset as above, using this reduced version of the family variable, and extracted all fitted parameters from the GLMM to form the predictive model. We did not include information on whether individuals were male or female, even though male and female moths can differ substantially in size in some species, because this information is not recorded in historical abundance datasets. Our model therefore used overall slope and intercept to predict body mass from forewing length for all moths, with a refined prediction for moths from the most speciose (and therefore data-rich) four families in our dataset.

### Testing model accuracy

To test the accuracy of this general predictive modelling approach when making predictions based on forewing length data from field guides, we estimated the body mass of each of the 94 moth species in our dataset from its expected forewing length (obtained by taking the midpoint between minimum and maximum forewing lengths given by field guides for micro-moths (Sterling & Parsons, 2012) and macro-moths (Waring & Townsend, 2017); Table S1), and used these estimates to calculate the estimated biomass of each mixed-species sample of moths (where one sample = all moths that were captured at the same site on the same day, across multiple traps; n = 44 samples). We compared between these estimates of biomass, and the empirically-measured biomass of the moths in question. We conducted this testing at both species- and sample-levels, because rare species from rare families are likely to have the least accurate predictions from our model, but may also have the least impact on the accuracy of sample-level predictions.

We first compared between measured and estimated biomass for the full set of 600 moths. At both species- and sample-level, we tested the relationship between measured and predicted biomass, using model II regressions with a Major Axis approach because neither biomass variable was dependent upon the other (Legendre & Legendre, 2012). Significance of relationships from random was tested using one-tailed permutation tests (with 100 permutations), and relationships were also compared to the desired y = x (i.e. estimated = measured) relationship by calculation of 95% confidence intervals around the estimated slope. The strength of the relationships between measured and estimated biomass at species- and sample-level was determined by model R^2^ values.

However, because in this case comparisons were not independent of the predictive model (i.e. model accuracy was tested with the same data that had been used to fit the model), we also used a resampling approach to further test the accuracy of our general predictive modelling approach. We split our full dataset 10,000 times into training and testing subsets. In each replicate, we randomly selected 480 individual moths (80 % of the 600 total individuals) without replacement to form a training subset, with the remaining 120 individuals forming an independent testing subset. We trained a model with the same structure as the full predictive model (above) on the training dataset, and from its parameters, extracted estimates of species- and sample-level biomass as above for the 120 moths included in the testing dataset. We tested the relationship between measured and predicted biomass for each replicate as above. Across the results of all 10,000 replicates (and at both species- and sample-levels), we then calculated the proportion of replicates for which measured and estimated biomass were significantly correlated, the mean and standard error of model R^2^ values, and the proportion of replicates for which the modelled relationship was significantly different from y = x.

Finally, we used a resampling approach to assess the influence of moth abundance (i.e. sample size) on prediction error. We randomly sampled sets of individuals (with replacement, from the full set of 600 measured individuals) at sample sizes between 10 and 1000 in steps of 10, taking 1000 replicates at each sample size for a total of 100,000 replicates. For each replicate sample, we calculated the measured biomass and the estimated biomass (based on the parameters of the final predictive model). We then calculated the prediction error for each sample as a proportion of the true biomass, normalizing by subtracting the known prediction error of 3.40 % in the full dataset (i.e. the total predicted biomass of all 600 moths was 3.40 % lower than their total measured biomass), such that:

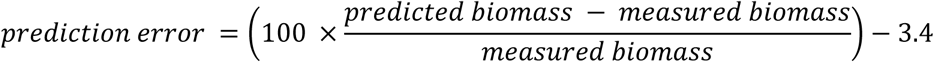

Grouping sample sizes into windows of 100, we calculated the mean, standard error and range of prediction errors observed across all replicates in each window.

All statistical analyses were conducted in R version 3.6.1 (R Core Team, 2019) using the following packages: *lme4* to fit and assess linear mixed-effects models (Bates et al., 2015); *lmodel2* to conduct model II regressions (Legendre, 2018); and *ggplot2* to plot figures (Wickham, 2016). All R scripts and data used in the analysis are archived online at Zenodo (doi: 10.5281/zenodo.3734274).

## Results

### Field sampling, identification and measurement of moths

We sampled 614 individual moths, of which 13 could not be confidently identified beyond family level. One micro-moth (*Narycia duplicella* [Goeze, 1783], Psychidae) could not be detected by our balance (and therefore weighed less than 0.005 mg). These 14 individual moths were excluded from further analyses. The remaining dataset contained exactly 600 individual moths, representing 94 species from 11 families (6.6 % of all species, or 13.7 % of macro-moth species, ever recorded in vice-county 61; Table S2). Among these moths, forewing lengths ranged from 7 mm (individuals of *Eudonia pallida* (Crambidae) and *Agapeta hamana* (Tortricidae)) to 40 mm (an individual of *Laothoe populi* (Sphingidae)) and dry body masses ranged from 1.1 mg (an individual of *Eupithecia tenuiata* (Geometridae)) to 753.2 mg (an individual of *Smerinthus ocellata* (Sphingidae)).

### Modelling forewing length – body mass relationship from empirical data

From the three candidate model structures described above, we selected the non-linear predictor (model ii) as the best-fitting model (BIC: 360.7, compared to 431.1 and 494.3 for models i and iii respectively). The natural logarithms of body mass and forewing length were significantly related to each other at both species and individual levels (Fig. 1), with variation among the 11 families in the slope and intercept of this relationship (individual level: X^2^ = 35.9, d.f. = 10, *P* < 0.001; marginal R^2^ = 0.819) revealing that interfamilial variation in body plan significantly influences the scaling of forewing length to body mass.

**Figure 1:**
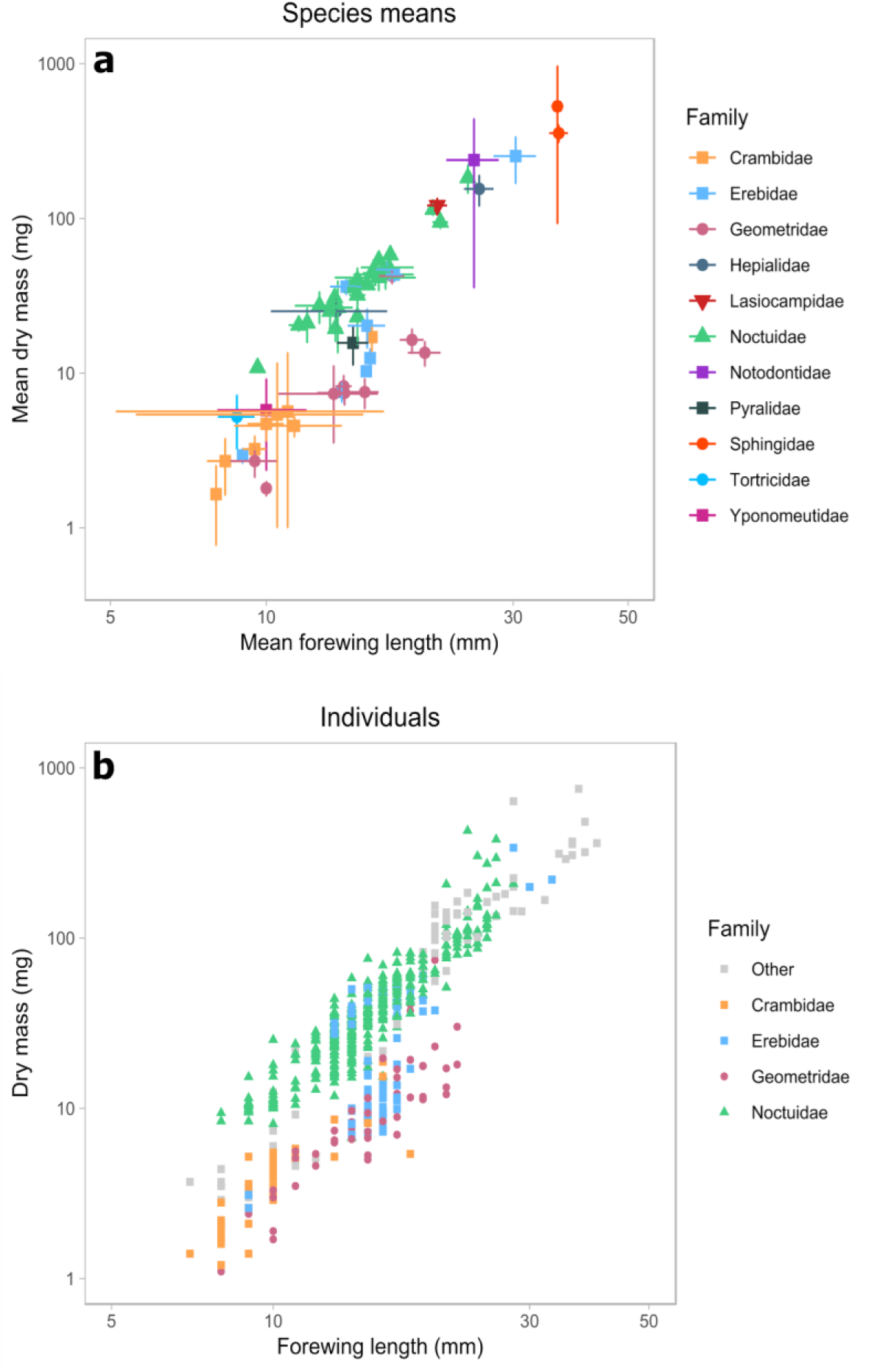
Relationship between forewing length (mm) and dry mass (mg). In the top panel, the mean forewing length and dry mass of each species sampled in the study is shown on logarithmic axes, with error bars showing standard errors and family indicated by the combination of point colour and shape. In panel (b), the forewing length and dry mass of every individual moth sampled in the study is shown on logarithmic axes, with the four most speciose families in our sample (Crambidae, Erebidae, Geometridae and Noctuidae) indicated as above by point colour and shape.

The significance of the model (and almost all of its explained variance) was retained when fitting the simplified model (in which seven families represented by < 5 species were grouped as ‘other’; X^2^ = 30.7, d.f. = 4, *P* < 0.001; marginal R^2^ = 0.812), resulting in a set of parameters from which body mass could be predicted based on forewing length (Table 1). All four families retained as independent levels (Crambidae, Erebidae, Geometridae and Noctuidae) had larger intercepts and shallower slopes than the overall prediction across the other families (Table 1). Thus we conclude that the non-linear model with simplified family variable has the greatest potential for estimating body mass.

**Table 1:** Parameters of the predictive model, extracted by fitting a GLMM with the fixed-effects structure: ln(body mass) ∼ ln(forewing length) × family, to data from 600 individual moths. The number of measured individuals and species on which each parameter estimate was based is given. Overall model parameters are given, including the X^2^ and *P-*values of a Likelihood Ratio Test of the model’s overall significance. Family-specific slope and intercept values are refinements to be added to the parameters for ‘other families’ (rather than taken in isolation). To predict body mass of a moth from its forewing length, these parameters should be applied to the following formula: ln(body mass) = (ln(forewing length) × (‘other families’ slope + family slope adjustment)) + (‘other families’ intercept + family intercept adjustment).

| Family adjustment | $n$ species ($n$ individuals) | $\chi^2$, d.f. ($P$) | Slope estimate (s.e.) | Intercept estimate (s.e.) |
| --- | --- | --- | --- | --- |
| Overall model | 94 (600); 11 families | 30.7, 4 (<0.001) | - | - |
| 'Other families' (no adjustment) | 15 (67); 7 families | - | 3.056 (0.180) | -5.016 (0.540) |
| Crambidae | 11 (38) | - | -0.904 (0.311) | 1.361 (0.813) |
| Erebidae | 10 (79) | - | -0.601 (0.360) | 1.294 (1.029) |
| Geometridae | 22 (52) | - | -0.492 (0.322) | 0.344 (0.891) |
| Noctuidae | 36 (364) | - | -1.297 (0.239) | 3.788 (0.694) |

### Testing model accuracy

We then used our best-fitting model to estimate body masses for all 94 species as described above, and compared between measured and estimated biomass for the full sample of 600 individual moths. We found that our estimates of biomass significantly correlated with measured biomass at both species- and sample-levels (Fig. 2), even though body mass varied widely both within and between species (within-species s.d. of body mass = 34.6 mg, between-species s.d. of body mass = 74.7 mg). At sample-level, the relationship between estimated and measured biomass was not significantly different from a 1:1 relationship (Table 2), with 91.5 % of variation explained. At species-level, estimated biomass explained 91.1 % of variation. The relationship was less steep than the expected 1:1 relationship (Table 2) with all moths included; however, the 1:1 relationship was recovered when we excluded the 34 smallest species from models (i.e. only included species weighing > 15 mg, n = 60 species). These results indicate that our predictive model may slightly overestimate the body mass of very small species of moths, but that this does not substantially bias estimates of sample-level biomass.

**Table 2:** Details of statistical models testing the relationships between measured biomass and estimated biomass at species- and sample-level for the final model. Relationships were tested using a model II regression, and significance was determined by a one-tailed permutation test with 100 permutations. The R^2^ of each model is also given, alongside the estimated intercept and slope of each model, with associated 95% confidence intervals.

| Level | Data subset | $n$ | Model $R^2$ | Model intercept<br>(95% CI) | Model slope<br>(95% CI) | $P$ |
| --- | --- | --- | --- | --- | --- | --- |
| Sample | Full dataset | 44 | 0.915 | 0.275<br>(-0.310 - 0.810) | 0.952<br>(0.865 - 1.047) | 0.010 |
| Species | Full dataset | 94 | 0.911 | 0.557<br>(0.382 - 0.723) | 0.874<br>(0.819 - 0.932) | 0.010 |
|  | Species weighing<br>> 15 mg<br>only | 60 | 0.823 | 0.168<br>(-0.311 - 0.595) | 0.964<br>(0.853 - 1.090) | 0.010 |

**Figure 2:**
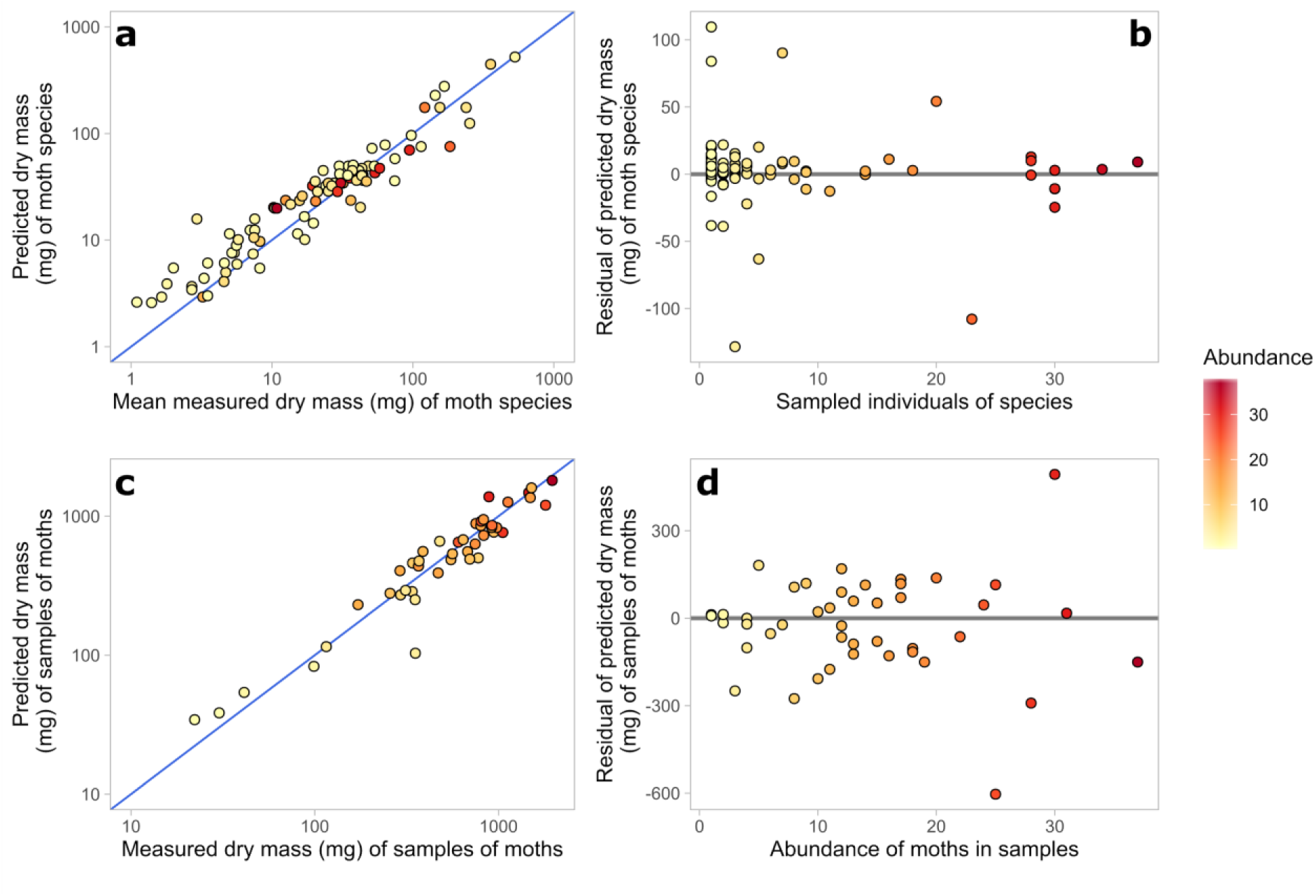
Accuracy of predicted biomass of moth species and samples of moths compared to the true, measured biomass. (a) Predicted dry mass of species (mg) is plotted against mean measured dry mass (mg); the 1:1 relationship is plotted as a blue line, and points are coloured by the number of individual moths from which the measured mean was calculated. (b) The absolute difference between mean measured dry mass and predicted dry mass of each moth species is plotted against the number of individuals from which the measured mean was calculated; a horizontal line is plotted at y = 0. (c) Predicted dry mass of samples (mg) is plotted against measured dry mass (mg); the 1:1 relationship is plotted as a blue line, and points are coloured by the number of individual moths contained in the sample. (d) The absolute difference between measured and predicted dry mass of each sample of moths is plotted against measured dry mass (mg); a horizontal line is plotted at y = 0.

To test whether this general predictive modelling approach can accurately estimate biomass beyond the sampled individuals and species, we split our data 10,000 times into random training (480 individuals in each case) and testing (120 individuals) subsets. We refitted our final model to the training subset in each case and predicted the body masses of individuals in each testing subset. We found again that our estimates of biomass significantly correlated with the measured biomass at both species- and sample-levels in 100% of replicates (Table 3). At sample-level, estimated biomass explained on average 88.4 % (± s.e. 0.07) of variation in measured biomass, and was not significantly different from a 1:1 relationship in 75.6 % of cases (Table 3), despite predictive models being built on a substantially reduced dataset compared to our final model. At species-level, estimated biomass explained 87.4 % (± s.e. 0.03) of variation in measured biomass, but the relationship was not significantly different from the expected 1:1 relationship in 19.5 % of cases (being significantly less steep than the expected relationship in the remaining 80.5 %: Table 3). As above, when we excluded the species weighing < 15 mg from testing subsets, the relationship was not significantly different from 1:1 in 81.3 % of cases (Table 3). These results indicate that predictions made using this approach are likely to remain accurate even when predicting beyond the training dataset.

**Table 3:** Details of bootstrap testing (over 10,000 replicates) of statistical models testing the relationships between measured biomass and estimated biomass at species- and sample-level. Each model was fitted to a training dataset consisting of 480 randomly-selected individuals, and tested on the remaining 120 individuals. Relationships were tested using a model II regression, and significance was determined by a one-tailed permutation test with 100 permutations. The R^2^ of each model was also taken, alongside the 95% confidence intervals for the estimated slope. Here, the number of replicates (/10,000) for which measured and estimated biomass were significantly related is given, as well as the number of replicates for which the 95% confidence intervals for the estimated slope contained 1 (i.e. y = x). The mean model R^2^ (and standard error) across all 10,000 replicates is also given. For tests of the slope’s relationship to 1, all models were re-tested with the species weighing > 15 mg excluded from the testing dataset.

| Level | Data subset | Bootstrap replicates | % models significant | % models slope not equal to 1 | Mean model $R^2$ (s.e.) |
| --- | --- | --- | --- | --- | --- |
| Sample | Full dataset | 10,000 | 100 | 24.4 | 0.8843<br>(0.0007) |
| Species | Full dataset | 10,000 | 100 | 80.5 | 0.8736<br>(0.0003) |
|  | Species weighing > 15 mg only | 10,000 | - | 19.5 | - |

Testing the influence of sample size on prediction error, we found that prediction error decreased initially as sample size increased, but remained relatively stable for samples larger than approximately 250 moths (Fig. S4). For samples of 10-100 moths, the standard error of prediction error was 0.13 (range −61.79 to 83.42 %), whereas for samples of 910-1000 moths, the prediction error was much less variable (s.e. 0.03, range −9.20 to 10.23 %). This indicates that sample-level estimates of biomass are especially accurate for samples containing > 250 moths.

## Discussion

Findings from our analyses show a strong relationship between forewing length and body mass in moths, which enables prediction (to an informative level of accuracy) of the biomass of samples of moths when such data are not available (e.g. because historical specimens have not been kept). Generating biomass data using this approach will provide an additional tool to ongoing investigation of the nature and consequences of changes in insect populations (Hallmann *et al*., 2017; Macgregor *et al*., 2019b; Didham *et al*., 2020b) using long-term recording datasets. It may also permit the inclusion of estimates of moth body mass in comparative studies and trait-based analyses, despite the general lack of empirical data of this nature (García-Barros, 2015). In particular, these data will facilitate studies of the relationships between biomass, abundance and community composition (Appendix S2), including important ecological questions such as: do biomass declines indicate a general decline in the abundance of the majority of species, a severe decline in the biomass of a few key species (e.g. Shortall *et al*., 2009), a shift in community composition towards smaller-bodied species, all of the above, or something else entirely?

### Evaluation of the predictive model’s current and future utility

Overall, the estimates of body mass calculated using the predictive model’s parameters performed relatively well during testing, with ∼ 90 % of variation in measured biomass explained by predicted biomass at both species- and sample-levels, and prediction error decreasing as sample size increased (Fig. S4). Therefore, using estimated body masses from the model (Table S1) to calculate the combined biomass of large samples of moths should yield accurate results. Our sampled dataset was only sufficiently data-rich to allow refined parameter estimates for four families (Crambidae, Erebidae, Geometridae and Noctuidae). Further improvement of the model’s accuracy may be possible by means of family-specific parameter estimates for other families, at sub-family level (to better account for within-family variation in body plan), or potentially even through a phylogenetic imputation approach (Penone *et al*., 2014). However, all of these approaches require the collection of additional data on body masses from a greater number of species from families which are less abundant in moth-trap samples than the dominant family, Noctuidae, and so collection of sufficient data using field sampling may be challenging. An alternative may be to use museum collections to measure individuals of a much wider range of species and families, where methods exist to account for the mass of entomological pins when taking such measurements (Gilbert, 2011). Taking such an approach might allow for more data to be collected even from rarely-trapped families (e.g. Sphingidae), or those which are speciose globally but have few (e.g. Saturniidae) or no (e.g. Hedylidae) species extant in Britain, thereby potentially allowing refined predictions to be made for many more families, potentially even at a global scale. Nevertheless, estimates of British moth biomass made using our approach (Macgregor *et al*., 2019b) revealed that 93.3% of total biomass is comprised of the three macro-moth families for which we made refined predictions (Erebidae, Geometridae and Noctuidae), so improving prediction accuracy for other families may have limited impact on the sample-level accuracy of the overall model.

One source of potential error when using published forewing lengths to estimate biomass is that 19% of individuals in our 2018 dataset had a measured forewing length which was outside the expected range given by field guides. Nevertheless, there was an overall correlation (R^2^ = 0.942) between the mean forewing length at species-level derived from our 2018 empirical measurements and the midpoint of the range of forewing lengths for each species, taken from the published field guides (Fig. S5). This suggests sufficient accuracy in our approach, particularly considering that our largest measured species had a forewing length 571 % larger than that of our smallest species. Similarly, the approaches we took to measuring forewing lengths (i.e. with analogue callipers and a ruler, to the nearest 1mm) and dry body masses (i.e. after one week of air-drying) mean that our dataset may not be fully comparable to datasets collected under other conditions or using other approaches (e.g. using digital callipers with higher resolution to measure forewing length, or measuring dry body mass after oven-drying). However, since all air-drying took place, and all measurements were taken, by the same person in the same laboratory over the same 6-week period (and air-drying for one week was shown to be sufficient for the mass of even the largest moths to stabilise: Fig. S2), these measurements are adequate to accurately establish the relative relationships between species for both forewing length and dry body mass. Therefore, our models can also be safely used to estimate relative change in moth biomass over time, or in space, assuming only that the average body mass of each individual species does not substantially change over the same scales.

An additional source of possible error in our models is sexual dimorphism in moths. Some moth species, including some sampled in our study (e.g. Drinker *Euthrix potatoria*; Lasiocampidae), exhibit substantial sexual dimorphism in wing length (Waring & Townsend, 2017) and in body mass (Allen *et al*., 2011). However, we did not quantify or adjust for sexual dimorphism in this study because long-term recording schemes rarely include information on sex of individual moths, even for dimorphic species, although the majority of such records are likely to be males (Altermatt *et al*., 2009). Therefore, for estimation of sample-level biomass, it will be of most use to provide a single average estimate of body mass per species, regardless of size dimorphism.

### Future research using our predictive model to study biomass change

Questions remain regarding temporal, spatial, and taxonomic variation in observed biomass declines (Shortall et al., 2009; Macgregor *et al*., 2019b; Didham *et al*., 2020b), the potential drivers of these declines (Grubisic *et al*., 2018; Komonen *et al*., 2019; Didham *et al*., 2020a), and the challenges of extrapolating across data types, geographic locations, and temporal and spatial scales (Thomas *et al*., 2019; Wagner, 2019; Didham *et al*., 2020b). Our study illustrates the power of predictive models of body mass to tackle these challenges. Applying these estimates in the same way to RIS datasets across the UK over longer time-scales, or to other long-term moth abundance datasets, such as the National Moth Recording Scheme or the Garden Moth Scheme (Fox et al., 2011; Bates et al., 2014), will facilitate investigation of declines over longer time-periods and broader geographical scales than has previously been feasible. Moreover, the same general approach could be used to estimate body mass of moths in other databases, including macro-moth recording schemes from other regions (e.g. the Noctua database; Groenendijk & Ellis, 2011) and micro-moths, which were incorporated into the NMRS in 2016. Combining with similar existing models for other insect families and invertebrate taxa (Sage, 1982; Sample *et al*., 1993; Sabo *et al*., 2002; Höfer & Ott, 2009) could facilitate comparison of biomass losses across multiple datasets and taxa at a global scale. However, a possibility that cannot be accounted for using this approach is change over time in the mean body mass of individual moths (Wu *et al*., 2019), e.g. in response to climate warming (Gardner *et al*., 2011).

Our approach may also be of use for conducting trait-based analyses of moths (e.g. van Langevelde *et al*., 2018), where it is important that trait data have high precision (Middleton-Welling *et al*., 2018). Our predictive model offers a means to estimate body mass reproducibly, potentially across multiple data sources, using a trait (forewing length) that is straightforward to measure using basic equipment, and therefore can be robustly applied to other datasets. Previous trait-based analyses have used forewing length as a proxy for body size, but we have shown that there is interfamilial variation in this relationship (Fig. 1), which can be incorporated by using our approach. However, an appropriate level of caution is advised before applying our specific estimates of body mass to systems where the moth fauna is markedly different from that of the UK (e.g. tropical forests). Under such circumstances, using a parallel approach to fit a new, regionally-specific predictive model may be the most robust approach.

## Conclusions

We have developed a predictive model to estimate the dry body mass of moths based on their forewing length, using it to generate body masses for all British species of macro-moth. The predictions of sample biomass made by our model correlated strongly with measured biomass of the same samples (R^2^ = 0.915), indicating that this approach provides a robust way to estimate the biomass of samples of moths identified to species level. Our approach unlocks new opportunities to study trends in moth biomass over time and over large geographic regions.

## Acknowledgements

This study was funded by the Department of Biology, University of York, through an internal grant awarded to C.J.M. and R.S.K. We are grateful to collaborators and staff who have contributed data from the Rothamsted Insect Survey, a BBSRC-supported National Capability. C.D.T., J.K.H. and C.J.M. were additionally supported by the Natural Environment Research Council (grant no. NE/N015797/1).

## Author contributions

The study was conceived by C.J.M. and C.D.T., and designed by those authors in discussion with J.K.H. Field and laboratory work was conducted by R.S.K., who also carried out the statistical analysis with C.J.M. The first draft of the manuscript was written by R.S.K. and C.J.M., and all authors contributed to subsequent revisions.

## Data Accessibility

All R scripts and data used in the analysis are archived online at Zenodo (doi: 10.5281/zenodo.3734274).

## Competing Interests

The authors declare no competing interests.

## Notes

#### Summary of Updates

Improved testing of model accuracy and precision. Case study analysis of historical dataset removed. Updated to include multiple recent relevant citations.

https://doi.org/10.5281/zenodo.3734274

