## Supplementary Information for "Unlocking the potential of historical abundance datasets to study biomass change in flying insects"

**Supporting Information**

**Appendix S1**: Detailed description of methods

*S1.1 Field sampling of moths*

Three sampling sites were selected, covering a range of habitat types with the aim of maximising the functional and taxonomic diversity of moths sampled (Fig. S1). On each sampling night, four traps were run from dusk until dawn (traps were turned on and off by a photocell switch), with two traps each placed at two of the sampling sites on a given night. Traps at the same sampling site were placed > 100 m apart and/or with intervening visual barriers, to reduce interference between traps. Sampling sites were rotated on a weekly basis so that each site was sampled for four weeks in total. Traps were emptied the following morning and captured moths were euthanised with ethyl acetate where possible, or by freezing at -20 °C for larger-bodied species. To limit impacts on local populations, we stopped collecting a given species once 30 individuals had been caught.

*S1.2 Quality control procedures for accuracy and precision of measurements*

To ensure that air-drying moths for a minimum of one week was sufficient to accurately measure their dry body mass, we measured the mass of a subset of 123 moths on a daily basis for nine consecutive days throughout the drying process (Fig. S2). We found that one week was sufficient for the mass of the majority of moths to stabilise, including the largest individuals (i.e. those with the smallest surface area: volume ratio, which might therefore be expected to dry most slowly).

To confirm that our measurements of each moth’s forewing length and dry body mass were precise, we took repeated measurements of five individuals each of two species at opposite ends of the spectrum of size: Middle-barred Minor *Oligia fasciuncula* (mean forewing length 9.6 mm ± 0.1) and Poplar Hawk-moth *Laothoe populi* (mean forewing length 36.7 mm ± 0.8). Across all individuals of both species, the mean coefficient of variation was 1.64 % per individual for forewing length (range 0 – 5.59 %) and 0.99 % per individual for dry body mass (range 0 – 4.77 %), indicating that most measurements were precise to within ± 6 %. Note that this is small compared to the 382 % difference between the forewing lengths of *O. fasciuncula* and *L. populi* respectively.

**Appendix S2**: Biomass-abundance relationships across samples

*S2.1 Methods*

We investigated the relationships between measured biomass, predicted biomass, abundance and species richness at sample-level across our three study sites in 2018. We used generalised linear mixed-effects models (GLMMs) to investigate pairwise relationships of abundance and species richness with each biomass variable, with site as a random effect in each model. We tested significance using a Likelihood Ratio Test. We also confirmed significance of relationships using model II regressions with a Major Axis approach, as above, because it was unclear which variable should be viewed as the independent variable (Legendre & Legendre, 2012).

*S2.2 Results*

Amongst the samples of moths collected in 2018 for this study, we found that measured sample biomass was significantly predicted by abundance and species richness across the three study sites (Fig. S6), with a stronger relationship to abundance, as might be expected (R^2^ = 0.566; Tables S3, S4). These results were qualitatively unchanged and quantitatively similar when the biomass of samples was estimated from the predictive model (Table S3). These results further illustrate that conclusions drawn from estimated biomass (rather than direct measurements of biomass) are likely to be robust.

**Figure S1**: Locations of the three field-sampling sites, shown within the University of York. Two trap locations were established per site, with sites indicated by symbol shape. Map data © 2018 Google.

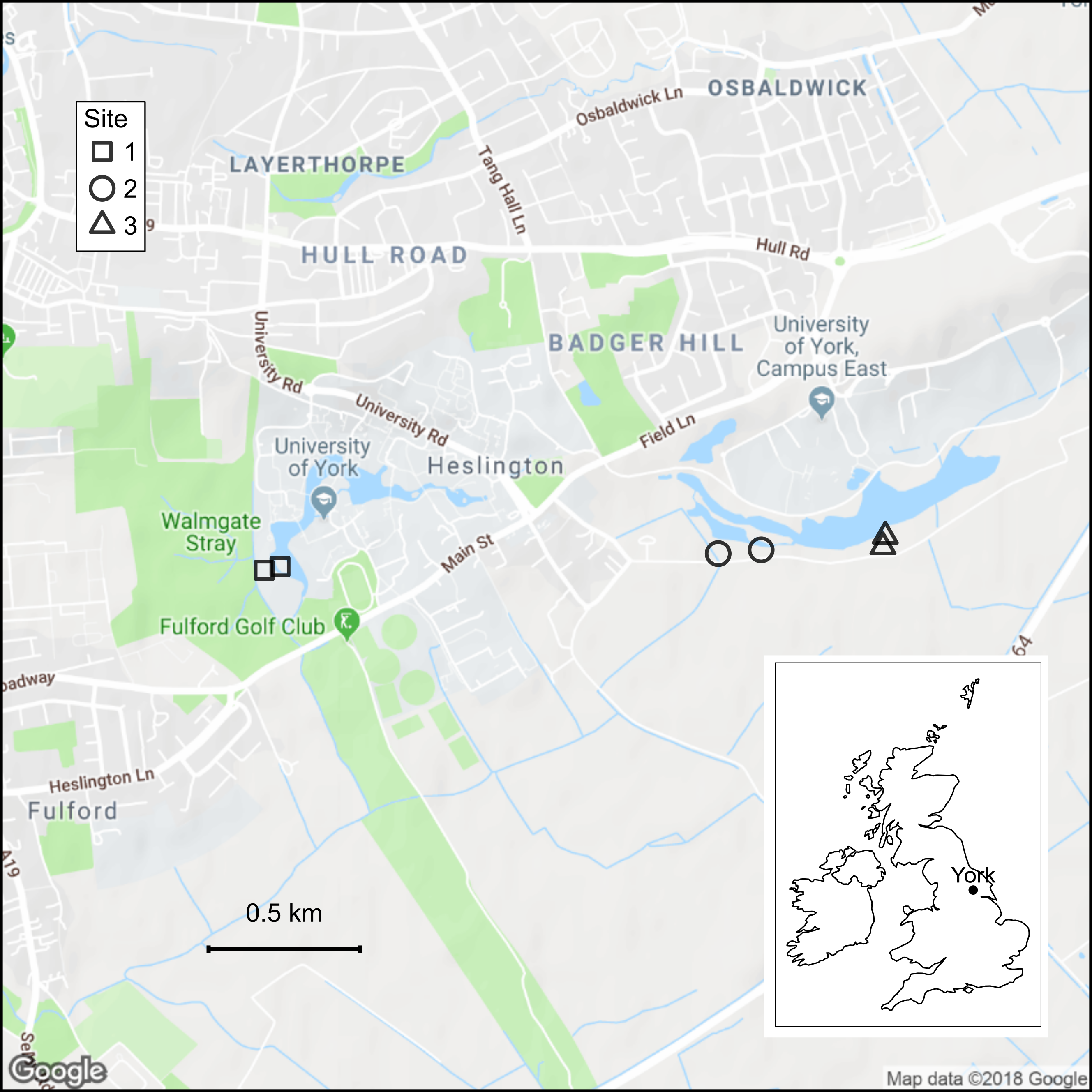

**Figure S2**: Daily decline and subsequent stabilisation in body mass for a subset of 123 moths measured over a 9-day period of air-drying. Moths were first measured (day 0) on the day after their initial capture, and were subsequently allowed to air-dry at room temperature. The mean daily mass of all measured moths is shown with 95% confidence intervals (dashed line), alongside the daily mass of the heaviest 20% of moths (excluding individuals not measured on day 8) (grey solid lines) and their mean daily mass with 95% confidence intervals (black solid line). Not every moth was measured on every day because no measurements were taken on weekends; therefore, apparent increases in mean masses were an artefact of the random subset of moths measured on each given day, rather than of actual increases in the mass of individual moths.

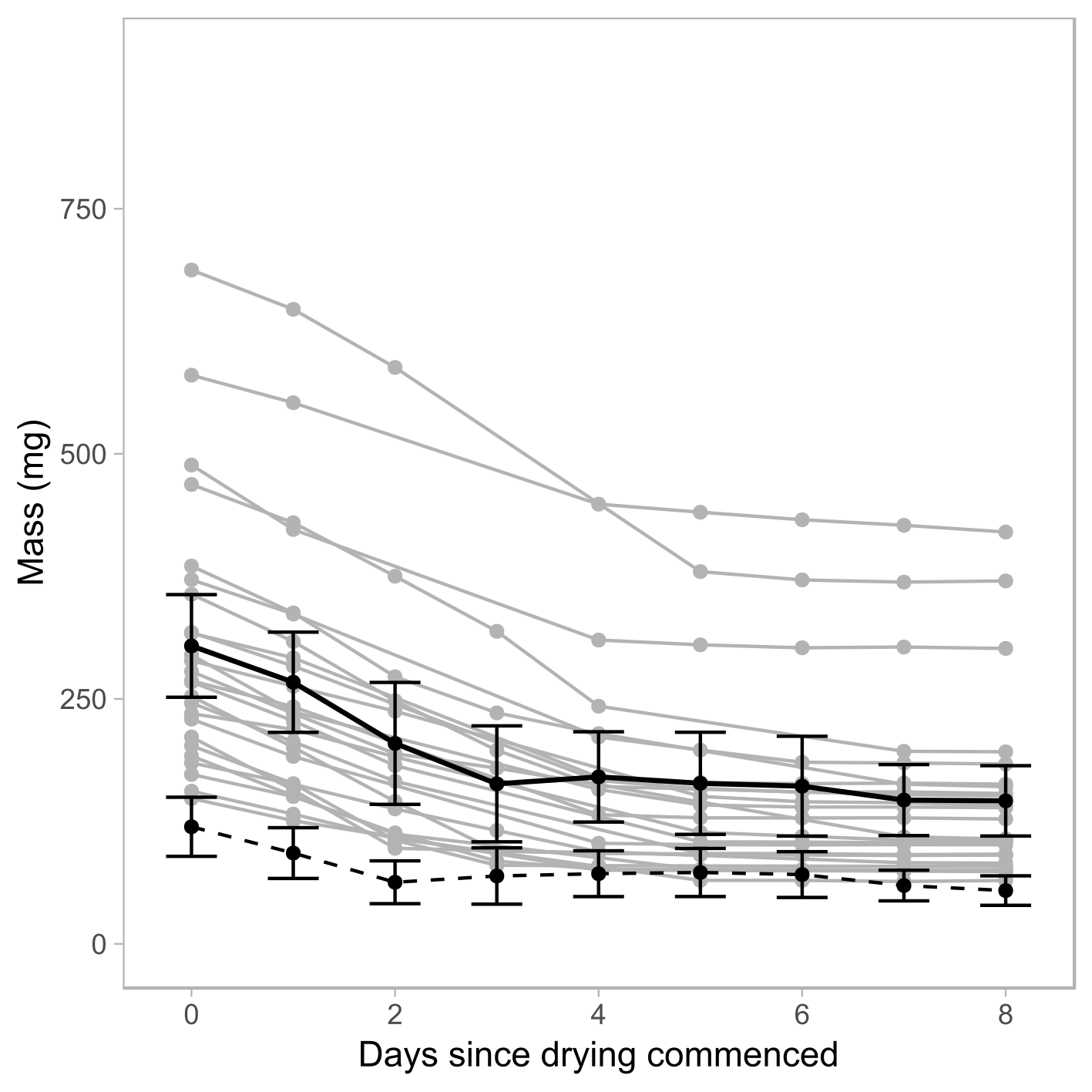

**Figure S3**: Predicted body masses of all British species of macro-moth plus micro-moths of the families Crambidae and Pyralidae, estimated using forewing length ranges extracted from Waring & Townsend (2017) and Sterling and Parsons (2012) respectively. Body mass was estimated using our predictive model at the midpoint between the minimum and maximum forewing lengths given by field guides.

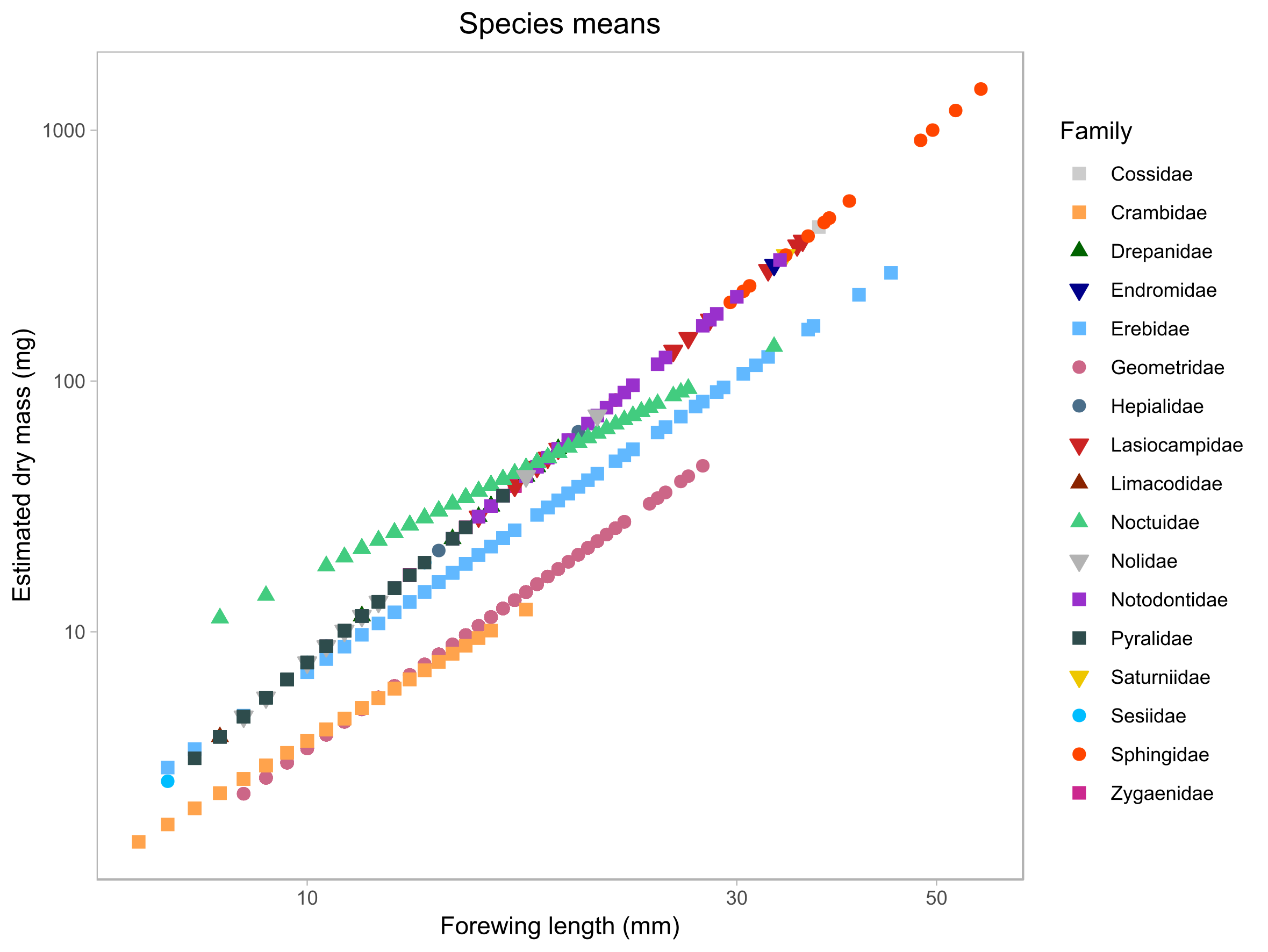

**Figure S4**: Effect of sample size on prediction error for estimated biomass, compared to measured biomass. Prediction error was calculated for 100,000 bootstrapped samples of individual moths sampled with replacement from the full dataset, each sample consisting of between 10 and 1000 moths. Prediction error for each sample is shown (grey points). Box-and-whisker plots summarise the data for sample-size windows of 100: boxes show median and quartiles, whiskers extend 1.5 × interquartile range beyond the boxes, and outliers are plotted.

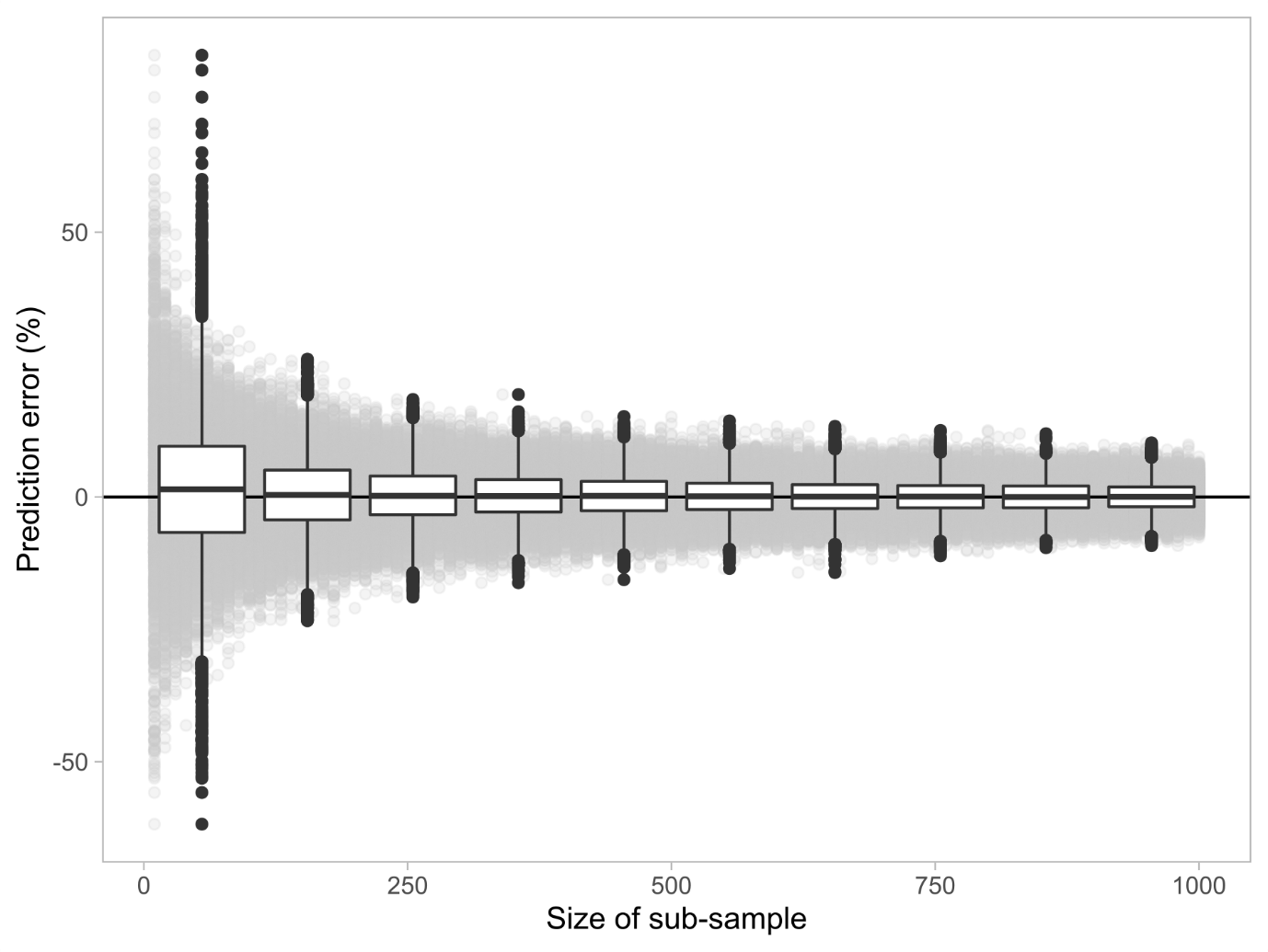

**Figure S5**: Measured forewing lengths of moths in the validation dataset correlated strongly and significantly with expected forewing lengths extracted from field guides, although the slope of this relationship was slightly less than 1. Each point represents a moth species, with the mean forewing length of all measured individuals on the y-axis. The expected y = x relationship is shown in blue, and the observed relationship fitted by a type II linear regression with a Major Axis approach is shown in black (slope: 0.937, 95% confidence interval: 0.890-0.986).

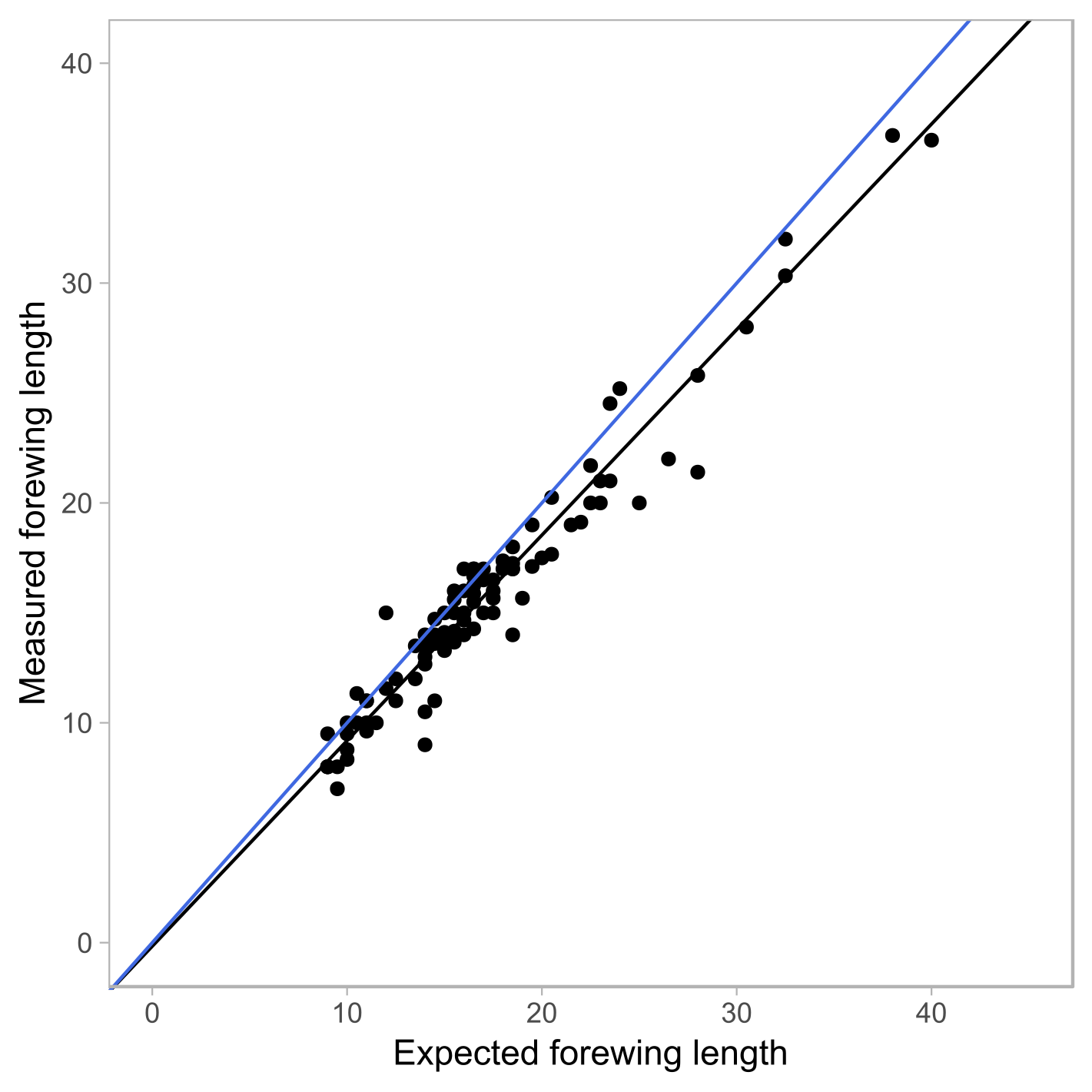

**Figure S6**: Relationship between sample biomass (mg) and abundance and species richness of moths, for (a,b) measured biomass of samples of moths captured in this study; and (c,d) predicted biomass of the same samples of moths captured in this study. Significant (*P* < 0.05) relationships are plotted as solid lines. Each point represents moths captured on a single night.

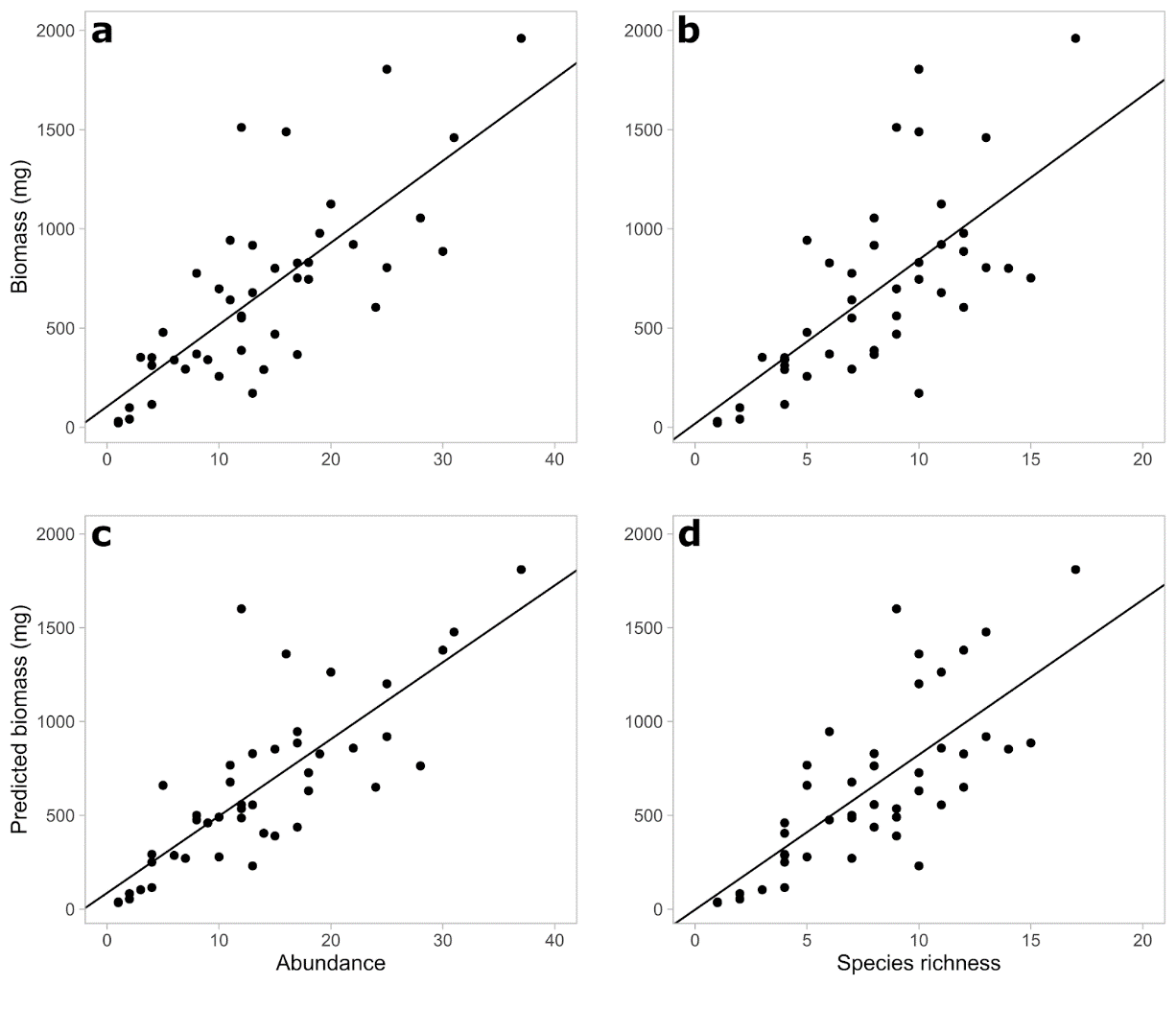

**Table S1**: Estimated dry body mass (mg) of all British species of macro-moth included in Waring and Townsend (2017) *Field Guide to the Moths of Great Britain and Ireland*. 3rd edition. London, UK: Bloomsbury Natural History, in addition to all members of the micro-moth families Crambidae and Pyralidae included in Sterling and Parsons (2012) *Field Guide to the Micro-moths of Great Britain and Ireland*. 1st edition. London, UK: British Wildlife Publishing. The natural logarithm of body mass (log_DRY_MASS) was estimated, with associated standard error (SE) using the predictive model (Table 1) at the midpoint (FOREWING_MED) between the minimum and maximum forewing lengths (FOREWING_LB and FOREWING_UB respectively) given in the field guide, and the estimate of body mass (PRED_DRY_MASS) therefore obtained by finding the exponential of this estimated value. [see attached spreadsheet]

**Table S2**: Species identity and abundance of 600 individual moths sampled during 2018 across 3 sites on the University of York campus, UK (Fig. S1), and identified to species level, for this study.

| Family | Common name | Binomial name | No. individuals sampled |
| --- | --- | --- | --- |
| Crambidae |  | *Agriphila straminella* | 14 |
|  |  | *Catoptria pinella* | 4 |
|  |  | *Chrysoteuchia culmella* | 6 |
|  |  | *Crambus perlella* | 2 |
|  |  | *Donacaula forficella* | 2 |
|  | Brown China-mark | *Elophila nymphaeata* | 1 |
|  |  | *Eudonia pallida* | 1 |
|  |  | *Eudonia truncicolella* | 3 |
|  | Ringed China-mark | *Parapoynx stratiotata* | 1 |
|  | Mother of Pearl | *Pleuroptya ruralis* | 2 |
|  |  | *Scoparia pyralella* | 2 |
| Erebidae | Garden Tiger | *Arctia caja* | 3 |
|  | Dingy Footman | *Eilema griseola* | 16 |
|  | Common Footman | *Eilema lurideola* | 28 |
|  | Yellow-tail | *Euproctis similis* | 3 |
|  | Beautiful Hook-tip | *Laspeyria flexula* | 2 |
|  | White Satin | *Leucoma salicis* | 1 |
|  | Ruby Tiger | *Phragmatobia fuliginosa* | 11 |
|  | Straw Dot | *Rivula sericealis* | 3 |
|  | White Ermine | *Spilosoma lubricipeda* | 3 |
|  | Buff Ermine | *Spilosoma luteum* | 9 |
| Geometridae | Peppered Moth | *Biston betularia* | 1 |
|  | Common White Wave | *Cabera pusaria* | 1 |
|  | Light Emerald | *Campaea margaritata* | 8 |
|  | Green Carpet | *Colostygia pectinataria* | 2 |
|  | Scalloped Oak | *Crocallis elinguaria* | 4 |
|  | Barred Straw | *Eulithis pyraliata* | 1 |
|  | Lime-speck Pug | *Eupithecia centaureata* | 1 |
|  | Slender Pug | *Eupithecia tenuiata* | 1 |
|  | Large Emerald | *Geometra papilionaria* | 1 |
|  | July Highflyer | *Hydriomena furcata* | 1 |
|  | May Highflyer | *Hydriomena impluviata* | 1 |
|  | Riband Wave | *Idaea aversata* | 9 |
|  | Small Fan-footed Wave | *Idaea biselata* | 2 |
|  | Single-dotted Wave | *Idaea dimidiata* | 2 |
|  | Clouded Border | *Lomaspilis marginata* | 1 |
|  | Green Pug | *Pasiphila rectangulata* | 1 |
|  | Willow Beauty | *Peribatodes rhomboidaria* | 4 |
|  | Shaded Broad-bar | *Scotopteryx chenopodiata* | 1 |
|  | Early Thorn | *Selenia dentaria* | 1 |
|  | Blood-vein | *Timandra comae* | 2 |
|  | Dark-barred Twin-spot Carpet | *Xanthorhoe ferrugata* | 1 |
|  | Silver-ground Carpet | *Xanthorhoe montanata* | 6 |
| Hepialidae | Ghost Moth | *Hepialus humuli* | 5 |
|  | Common Swift | *Hepialus lupulinus* | 3 |
| Lasiocampidae | Drinker | *Euthrix potatoria* | 20 |
|  | Oak Eggar | *Lasiocampa quercus* | 1 |
| Noctuidae | Poplar Grey | *Acronicta megacephala* | 4 |
|  | Dark Dagger | *Acronicta tridens* | 1 |
|  | Heart & Dart | *Agrotis exclamationis* | 30 |
|  | Shuttle-shaped Dart | *Agrotis puta* | 3 |
|  | Turnip Moth | *Agrotis segetum* | 1 |
|  | Copper Underwing | *Amphipyra pyramidea* | 2 |
|  | Mouse Moth | *Amphipyra tragopoginis* | 1 |
|  | Dark Arches | *Apamea monoglypha* | 30 |
|  | Dusky Brocade | *Apamea remissa* | 3 |
|  | Rustic Shoulder-knot | *Apamea sordens* | 2 |
|  | Silver Y | *Autographa gamma* | 1 |
|  | Beautiful Golden Y | *Autographa pulchrina* | 1 |
|  | The Flame | *Axylia putris* | 1 |
|  | Mottled Rustic | *Caradrina morpheus* | 1 |
|  | Treble Lines | *Charanyca trigrammica* | 1 |
|  | Dun-bar | *Cosmia trapezina* | 28 |
|  | Burnished Brass | *Diachrysia chrysitis* | 2 |
|  | Ingrailed Clay | *Diarsia mendica* | 14 |
|  | Dusky Sallow | *Eremobia ochroleuca* | 7 |
|  | Lychnis | *Hadena bicruris* | 2 |
|  | Uncertain | *Hoplodrina alsines* | 34 |
|  | Rustic | *Hoplodrina blanda* | 4 |
|  | Bright-line Brown-eye | *Lacanobia oleracea* | 3 |
|  | Common Rustic | *Mesapamea secalis/didyma* | 28 |
|  | Cloaked Minor | *Mesoligia furuncula* | 1 |
|  | Clay | *Mythimna (Hyphilare) ferrago* | 8 |
|  | Shoulder-striped Wainscot | *Mythimna comma* | 3 |
|  | Smoky Wainscot | *Mythimna impura* | 30 |
|  | Common Wainscot | *Mythimna pallens* | 5 |
|  | Large Yellow Underwing | *Noctua pronuba* | 23 |
|  | Flame Shoulder | *Ochropleura plecta* | 3 |
|  | Middle-barred Minor | *Oligia fasciuncula* | 37 |
|  | Marbled Minor | *Oligia strigilis* | 18 |
|  | Angle Shades | *Phlogophora meticulosa* | 1 |
|  | Setaceous Hebrew Character | *Xestia c-nigrum* | 1 |
|  | Double Square-spot | *Xestia triangulum* | 30 |
| Notodontidae | Buff-tip | *Phalera bucephala* | 5 |
|  | Lesser Swallow Prominent | *Pheosia gnoma* | 1 |
|  | Pale Prominent | *Pterostoma palpina* | 1 |
|  | Coxcomb Prominent | *Ptilodon capucina* | 1 |
| Pyralidae | Lesser Wax Moth | *Achroia grisella* | 1 |
|  | Bee Moth | *Aphomia sociella* | 7 |
| Sphingidae | Elephant Hawk-moth | *Deilephila elpenor* | 1 |
|  | Poplar Hawk-moth | *Laothoe populi* | 7 |
|  | Eyed Hawk-moth | *Smerinthus ocellata* | 2 |
| Tortricidae |  | *Agapeta hamana* | 9 |
| Yponomeutidae | Bird-cherry Ermine | *Yponomeuta evonymella* | 3 |

**Table S3**: Details of statistical models testing the relationships of sample biomass to abundance and species richness, using both directly measured biomass of samples of moths captured in this study and estimated biomass of the same samples (Fig. S6). Relationships were tested by fitting a GLMM and testing it using a Likelihood Ratio Test, so the test statistic shown is χ^2^. The marginal R^2^ of each model is also given, alongside the effect size, which represents the change in biomass (mg) for each unit of change in the explanatory variable (i.e. 1 individual or species).

| Biomass measure | Explanatory variable | *n* | Model R^2^ | Effect size (s.e.) | χ^2^ / *F* (*P*) |
| --- | --- | --- | --- | --- | --- |
| Measured biomass | Abundance | 44 | 0.566 | 41.3 (5.2) | 39.2 (<0.001) |
|  | Species richness | 44 | 0.474 | 82.7 (13.0) | 29.7 (<0.001) |
| Predicted biomass | Abundance | 44 | 0.614 | 41.0 (4.4) | 47.7 (<0.001) |
|  | Species richness | 44 | 0.532 | 82.6 (11.2) | 36.2 (<0.001) |

**Table S4**: Details of model II regressions re-testing the relationships of sample biomass to abundance and species richness, using both directly measured biomass of samples of moths captured in this study and estimated biomass of the same samples (Fig. S6). Relationships were tested using model II regressions with a Major Axis approach, and significance was determined by means of one-tailed permutation tests with 100 permutations. The R^2^ of each model is also given, alongside the effect size (and associated 95% confidence interval), which represents the change in biomass (mg) for each unit of change in the explanatory variable (i.e. 1 individual or species).

| Biomass measure | Explanatory variable | *n* | Model R^2^ | Effect size (95% CI) | *P* |
| --- | --- | --- | --- | --- | --- |
| Measured biomass | Abundance | 44 | 0.580 | 70.5 (55.8 - 96.0) | 0.010 |
|  | Species richness | 44 | 0.491 | 170.8 (129.7 - 250.0) | 0.010 |
| Predicted biomass | Abundance | 44 | 0.635 | 63.8 (51.6 - 83.5) | 0.010 |
|  | Species richness | 44 | 0.559 | 151.7 (118.8 - 209.7) | 0.010 |
